# Geospatial HIV-1 subtype C gp120 sequence diversity and its predicted impact on broadly neutralizing antibody sensitivity

**DOI:** 10.1101/2020.09.09.289132

**Authors:** Jyoti Sutar, Suprit Deshpande, Ranajoy Mullick, Nitin Hingankar, Vainav Patel, Jayanta Bhattacharya

## Abstract

Evolving diversity in globally circulating HIV-1 subtypes presents formidable challenge in defining and developing neutralizing antibodies for prevention and treatment. HIV-1 subtype C is responsible for majority of global HIV-1 infections. Broadly neutralizing antibodies (bnAbs) capable of neutralizing distinct HIV-1 subtypes by targeting conserved vulnerable epitopes on viral envelope protein (Env) are being considered as promising antiviral agents for prevention and treatment. In the present study, we examined the diversity in genetic signatures and attributes that differentiate region-specific global HIV-1 subtype C *gp120* sequences associated with virus neutralization outcomes to key bnAbs having distinct epitope specificities. A total of 1814 full length HIV-1 subtype C gp120 sequence from 37 countries were retrieved from Los Alamos National Laboratory HIV database (www.hiv.lanl.gov). The amino acid sequences were assessed for their phylogenetic association, variable loop lengths and prevalence of potential N-linked glycosylation sites (pNLGS). Responses of these sequences to bnAbs were predicted with a machine learning algorithm ‘bNAb-ReP’ and compared with those reported in the CATNAP database. Phylogenetically, sequences from Asian countries including India clustered together however differed significantly when compared with pan African subtype C sequences. Variable loop lengths within Indian and African clusters were distinct from each other, specifically V1, V2 and V4 loops. Furthermore, V1V2 and V2 alone sequences were also found to vary significantly in their charges. Pairwise analyses at each of the 25 pNLG sites indicated distinct country specific profiles. Highly significant differences (p<0.001***) were observed in prevalence of four pNLGS (N130, N295, N392 and N448) between South Africa and India, having most disease burden associated with subtype C. Our findings highlight that the distinctly evolving clusters within global intra-subtype C *gp120* sequences are likely to influence the disparate region-specific sensitivity of circulating HIV-1 subtype C to bnAbs.

## Introduction

The extraordinary diversity of *env* targeting neutralizing antibodies is a barrier to achieving the desired vaccine-induced and antibody-mediated protection. The evolving antigenic diversity in global and region-specific circulating HIV-1 subtypes is complex which not only poses significant roadblock to developing preventive vaccine but also poses challenge in neutralizing antibody mediated prophylaxis and treatment. Broadly neutralizing antibodies (bnAbs) act solely on the HIV-1 envelope glycoprotein (Env) in neutralizing genetically distinct HIV-1 subtypes. Till date a number of bnAbs have been discovered from elite neutralizers and few of them have been found to be prevent acquisition as well as significantly reducing the plasma viral loads, when tested both in animal models and humans (Escolano et al. 2017; R. Kumar et al. 2018a; Mendoza et al. 2018), thus justifying their importance as products for prevention and treatment. Some of the bnAbs that are currently being evaluated through human clinical trials should provide additional possibilities for prevention (Sok and Burton 2016, 2018), such as their extent, in addition to virus neutralization, in persistent viral clearance (Caskey et al. 2019; Nishimura and Martin 2017) and eliminating HIV-1 infected cells (Caskey et al. 2015; Caskey et al. 2017; R. Kumar et al. 2018a). While some single bnAbs have been found to show significant breadth across subtypes, combination of bnAbs with distinct epitope-specificity is believed to provide the most effective response against globally diverse HIV-1 subtypes and also would likely prevent the development of antibody-escape variants (R. Kumar et al. 2018a). The trimeric Env glycoproteins on the virus surface are the most diverse of all proteins encoded by HIV-1; which differs by greater than 20% of amino acids between matched subtype (Gnanakaran et al. 2007; Han et al. 2020; Hraber et al. 2014; Korber et al. 2001; Lynch et al. 2009) and which continues to diversify at a population level (Bouvin-Pley et al. 2013; Bunnik et al. 2010; DeLeon et al. 2017; Hraber et al. 2014). This has been substantiated by observation that several epitopes that are targeted by different bnAbs those have been found to vary over time (DeLeon et al. 2017). While bnAbs target both surface gp120 and membrane proximal external region (MPER) of gp41, gp120 exhibits extraordinary sequence divergence compared to that of gp41 MPER and variation in this region is believed to represent distinct genetic subtypes, or clades, which are prevalent in distinct geographic regions (Buonaguro et al. 2007; Taylor et al. 2008). Although bnAbs isolated from individuals infected with one particular subtype are generally effective at neutralizing viruses belonging to other subtypes, antibody potency is often found to be correlated with matched subtypes as described elsewhere (Binley and Burton. 2004; Bures et al. 2002; Hraber et al. 2014; Kulkarni et al. 2009; Li et al. 2006; Seaman et al. 2010). Moreover, diversity has been found to have an impact even within matched subtype, as demonstrated by the fact that the subtype-matched neutralization advantage was more apparent in regions with distinct viral diversities (Hraber et al. 2014).

HIV-1 subtype C accounts for approximately half of the global infections (Novitsky et al. 2002), which predominates in India and South Africa. Yet, robust, comparative *env* sequence diversity and evolution analysis, critical for bnAb based intervention strategies in these regions is severely lacking (Sutar et al. 2019). While recent development of bnAbs has considerably improved our knowledge on conserved epitopes that they target, and Env structure associated with broad and potent virus neutralizing antibodies, a greater understanding of the antigenic diversity of global HIV-1 subtype C Env would facilitate understanding potential bnAb combination that could potentially overcome the intra-clade C diversity. In the present study, we examined the variation in signature sequences, loop length, N-linked glycosylation and key epitopes within the existing globally circulating HIV-1 subtype C *gp120* targeted by potent bnAbs and predicted their potential impact on virus neutralization.

## Results

### Evidence of phylogenetic divergences of globally circulating HIV-1 subtype C*gp120* sequences

Previous studies have demonstrated that HIV-1 subtype C is the most abundant globally circulating subtype responsible for approximately 46.6% of the global HIV infections (Hemelaar et al. 2019), In the present study, we retrieved from HIV database (www.hiv.lanl.gov) a total of 23750 sequences covering the complete *gp120* gene (HXB2 coordinates: 6225-7758). To mitigate the intra-individual quasispecies bias, we applied ‘one sequence/individual filter’ retaining 1927 full length sequences. As shown in Table 1, there is uneven distribution of *gp120* sequences from different countries. Of the 37 countries analyzed, South Africa (ZA) was found to contribute in the database (www.hiv.lanl.gov) more than 1000 sequences, while Malawi (MW), Botswana (BW) and Zambia (ZM) contributed between 100 to 1000 sequences. Interestingly, complete *gp120* sequences were found from only 84 unique individuals from India (IN), while all other countries contributed approximately 50 or less sequences. Upon removal of identical sequences as well as those harbouring internal stop codons, a total of 1814 full length HIV-1C gp120 protein sequences were retained for further analysis. We next analyzed the phylogenetic properties of the subtype C *gp120* sequences that uniquely represent globally circulating subtype C across different geography. Average number of amino acid differences per site between sequences grouped by source country were estimated by MEGA (S. Kumar et al. 2018b). Sequences from Asian Countries (IN, China (CN) and Nepal (NP)) were observed to be closer to each other (0.183-0.208) compared to African countries (ZA and MW: 0.207-0.234). This trend continued in the maximum likelihood tree constructed as indicated in Figure 1A, wherein sequences from Asian countries (IN, CN and NP) were observed to form a unique sub-cluster. To further validate these observations statistically, a subset of sequences (N=251) forming the aforementioned diverging node were reanalysed with 1000 ultrafast bootstrap replicates and SH-aLRT. As indicated in figure 1B, sequences from Asian countries clustered together with 97% bootstrap and 92.1% SH-aLRT support. They also clustered distinctly from African sequences supported by 84.5% SH-aLRT and 79% ultrafast bootstrap replicates.

**Table 1:**
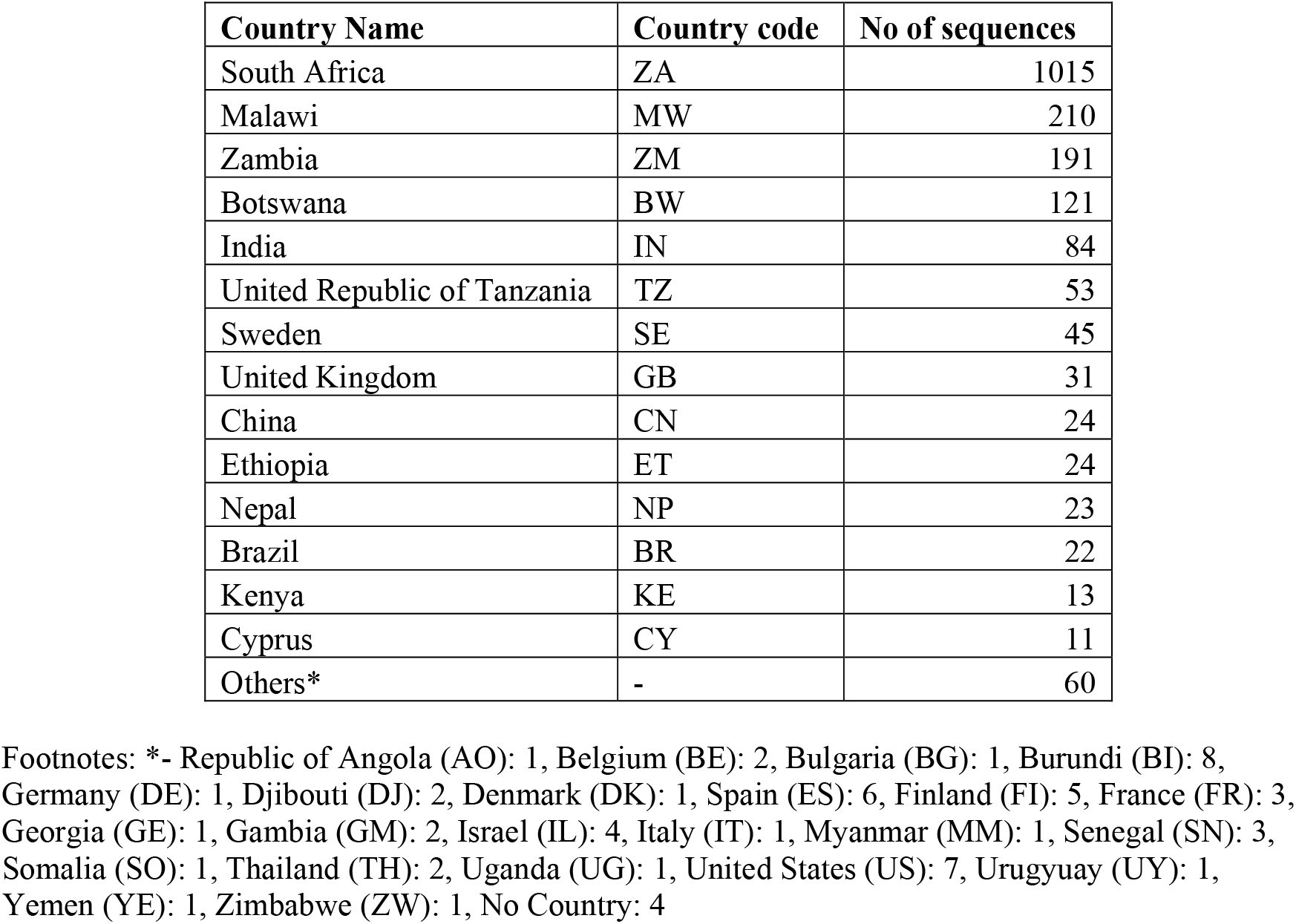
Details of Country-wise HIV-1 subtype C sequences retrieved from LANL-HIV database.

**Figure 1:**
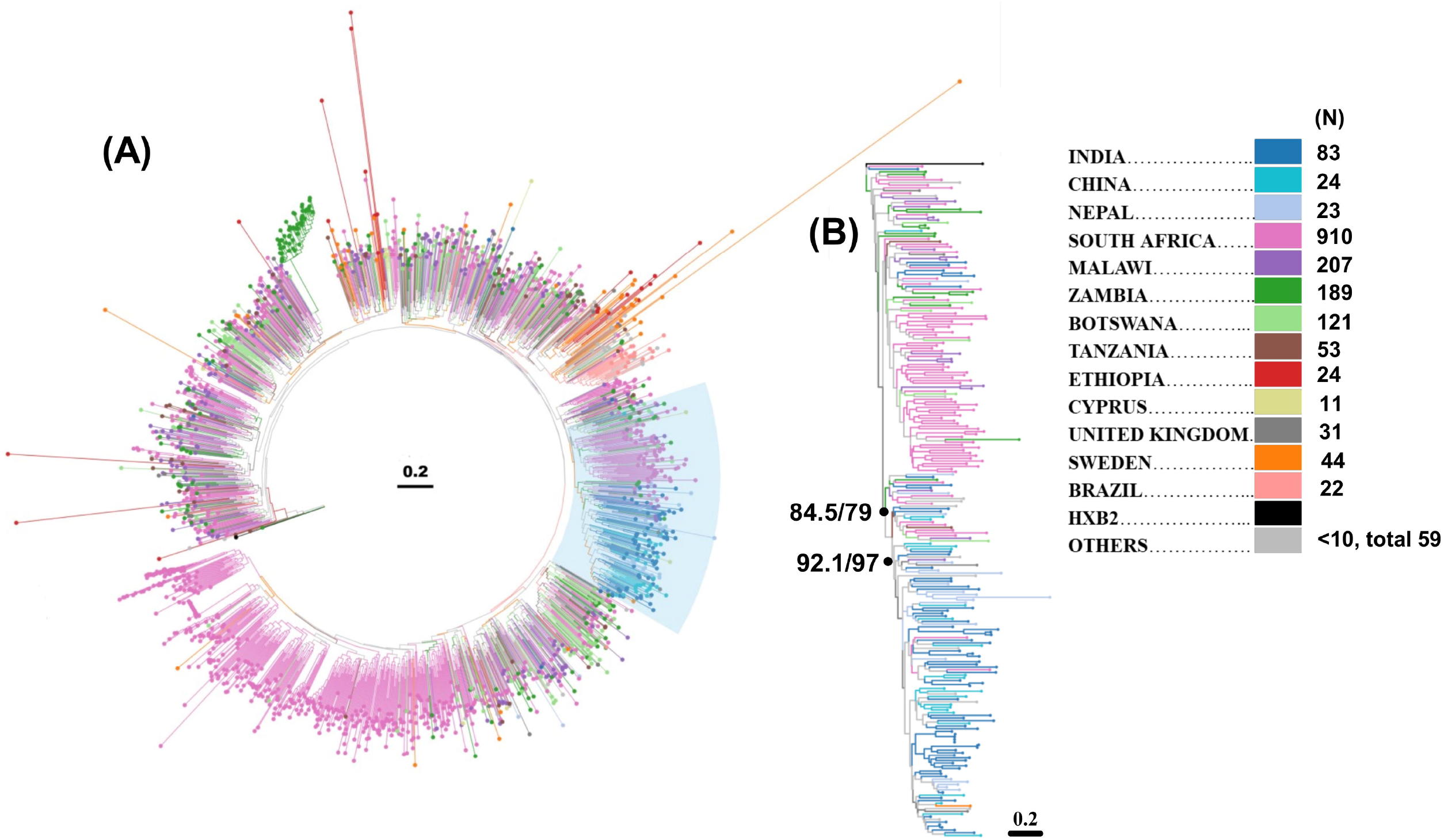
Phylogenetic Analysis of HIV-1 subtype C gp120 amino acid sequence. **A.** A maximum likelihood tree depicting phylogenetic association of 1837 HIV-1 subtype C amino acid sequences depicted radially. The legend describes the country codes as well as the sequence distribution among the countries. **B.** A maximum likelihood subtree detailing phylogenetic association between sequences from South Africa and Those from India, China and Nepal. The black dots indicate nodes with corresponding SH-aLRT and bootstrap values.

### Geography based subtype C*gp120* loop length and charge variation

Changes in the lengths of variable loops with gp120 has been previously documented to be an evasion mechanism of HIV-1 to escape neutralizing antibody driven humoral responses (Deshpande et al. 2016; Ringe et al. 2012; van Gils et al. 2011). Therefore, we next assessed lengths of variable regions between sequences from different countries. We included sequences from six countries in this analysis that could contribute more than 50 sequences each (Figure 2A). Comparison of variable region loop (V1-V5) lengths with Kruskal Wallis non-parametric test indicated several statistically significant differences between sequences from diverse countries. For the V1 loop, sequences from UK (GB) had significantly lower length (median:16) compared to all other countries (median range: 23-27). This trend continued in V2 loop with lower length in UK sequences (median: 34) compared to rest of the countries (median range: 42-48). Additionally, sequences from BW and NP were observed to be significantly distinct. However, this was an effect of outlier values and minimal standard deviation in the loop lengths respectively. These differences remained consistent when V1V2 loop were taken together indicating no loop length compensation. The V3 loop length was observed to be highly conserved across all the countries with some outlier values in South Africa. Interestingly, V4 loop length was observed to be most variable among the variable domains. Sequences from UK were observed to have significantly shorter V4 loop (Median: 13) compared to rest of the countries (Median range: 27-34). Of note, sequences from India (IN) had longer V4 loop (Median: 31) than those from South Africa (Median: 27). Sequences from African countries (BW, ET, KE, MW, TZ, ZA and ZM) had shorter V4 loop length (Median range: 27-30) compared to the rest of the countries (Median range: 29-34) except UK. Similar to the V3 loop, V5 loop length was conserved across all the countries under consideration (Median: 11) except UK (median: 6). Overall, length of the gp120 protein was observed to be shortest in sequences from GB (Median: 462) and longest in those reported from NP (Median: 518). Consistent with the V4 loop observations, gp120 length was observed to be lower (Median range: 503-511) in countries from Africa (BW, ET, KE, MW, TZ, ZA and ZM) compared to rest of the countries (Median range: 507-518) except UK. A subset of six countries with more than 50 sequences (IN, ZA, MW, TZ and BW) was further selected for assessment of sequence charge distribution (Figure 2B). V1, V1-hypervariable region, V2, V2 hypervariable region as well as V1V2 cumulatively were found to have significantly different distributions. Of note, V1V2 hypervariable region charge was distinct in India as compared to South Africa.

**Figure 2:**
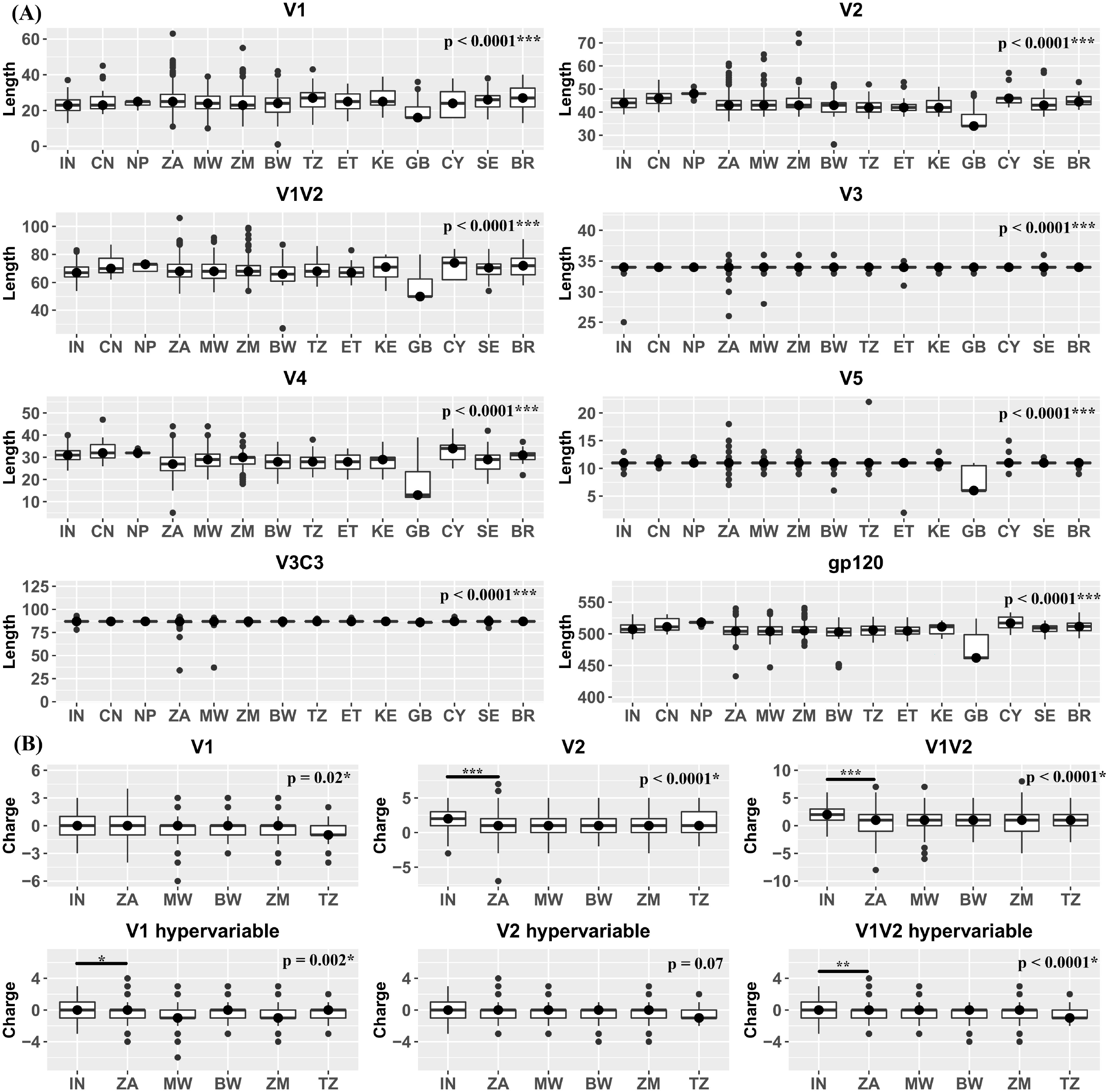
Assessment of Variable region characteristics. **A.** variable region length: gp120 variable region (V1, V2, V1+V2, V4, V5) as well as entire gp120 lengths have been plotted on the Y axis against the Countries of origin indicated on the X axis. **B.** Variable region charge: gp120 variable region Charges for V1, V2 and V1+V2 as well as hypervariable regions within them have been plotted on the Y axis against the Countries of origin indicated on the X axis. P values have been indicated following a statistical analysis by Kruskal-Wallis test followed by Dunn’s multiple comparison.

### Comparison of abundance of potential N linked glycosylation sites (pNLGs)

HIV-1 gp120 is a heavily glycosylated protein with host derived N-linked glycans making up ~50% of its total mass. These glycans play an important role in ensuring viral infectivity as well as evading neutralizing antibodies (Doores et al. 2015) emphasizing the importance of their assessment. In the present study, proportion of pNLGs were compared between sequences from the 14 aforementioned countries. pNLGs ranging from 11 to 33 were observed in the sequences. There were no overall significant differences between the number of average pNLGs between countries (Kruskal Wallis test, p>0.05). Out of a total of 3003 pairwise analyses performed with Fisher’s exact test at each NLG position between each of the countries, 342 combinations were found to be statistically significant (Fisher’s exact test, p<0.01*) involving 31 of the 33 pNLG sites. The two invariable sites were N140 (abundance: 17-33%) and N301 (abundance: 89-100%). N140, a position in the V1 hypervariable region is perhaps a displaced glycosylation position from N139/141 due to insertion/deletion, while N301 is an important glycosylation site that plays critical role in various envelope functions including membrane fusion and thus is highly conserved (Ogert et al. 2001). Figure 3 represents abundance at of the 25 well characterized pNLG sites denoted along with their functional domains. As apparent in figure 4 as well as observed in statistical comparisons, there was no significant difference between glycosylation profiles at any of the sites between India, China and Nepal. Similar observations were also made for a cluster of African countries South Africa, Tanzania, Kenya and Malawi. Furthermore, sequences from Ethiopia and Sweden were also observed to have similar NLG profiles. Upon comparison of sequences from India and south Africa, highly significant differences (Fisher’s exact test, p <0.001***) were observed in 4 pNLG sites (N130, N295, N392 and N448), which are present in C1, C2, V4 and C4 domains respectively. These sites are important for integrity of the ‘mannose patch’ and interaction with several bnAbs such as 2G12, VRC-PG05, PGT135 and PGT151 (Doores et al. 2015).

**Figure 3:**
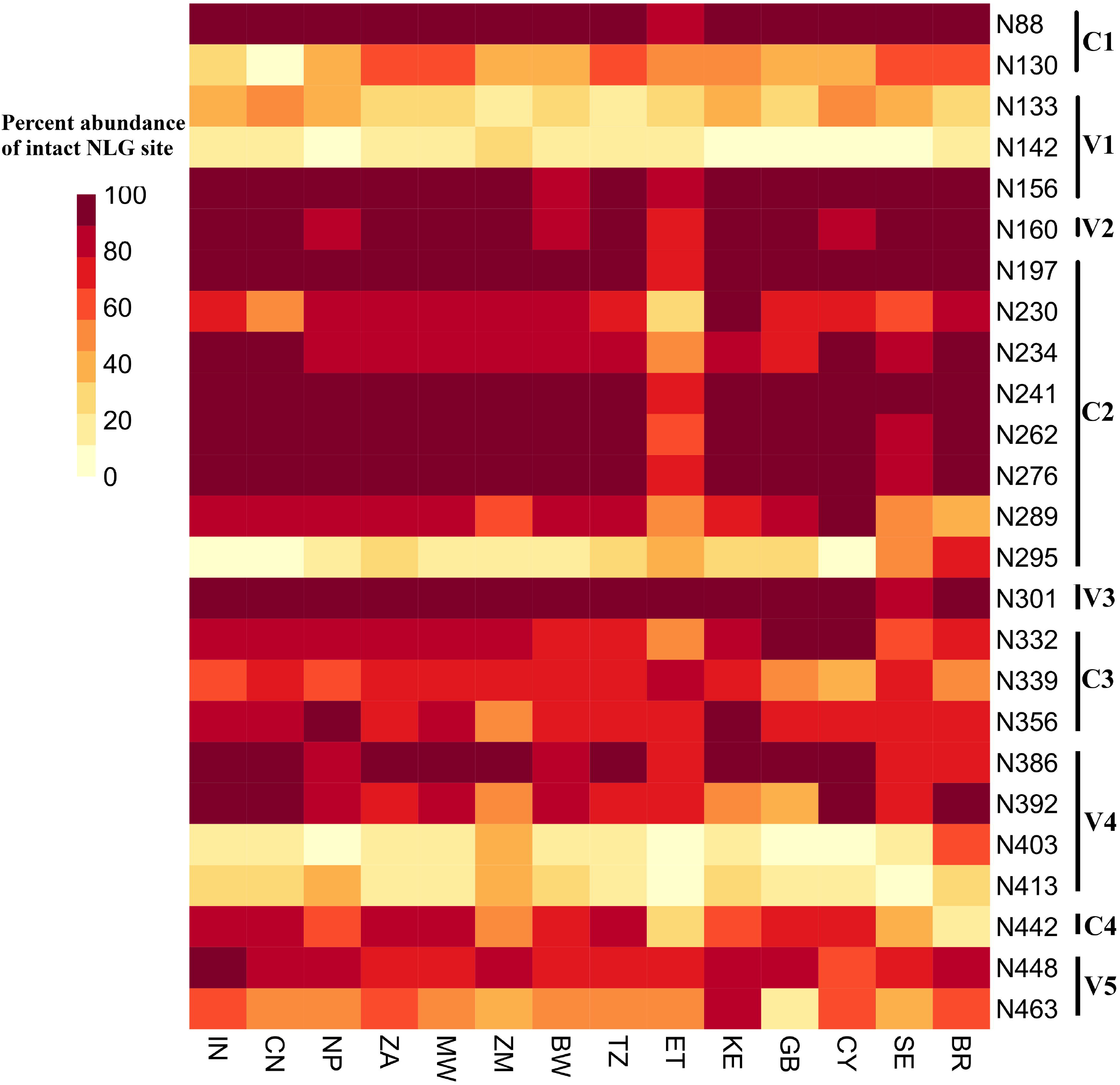
Assessment of potential N-linked glycosylation sites. A heatmap comparison of abundance of pNLG sites plotted on Y axis against countries plotted on X axis. Each pixel represents 1 pNLG site data from 1 country. Specific domains of pNLGs have been indicated along the Y axis. Color key represents correlation of color intensity with abundance of pNLGs ranging from 0 to 100.

**Figure 4:**
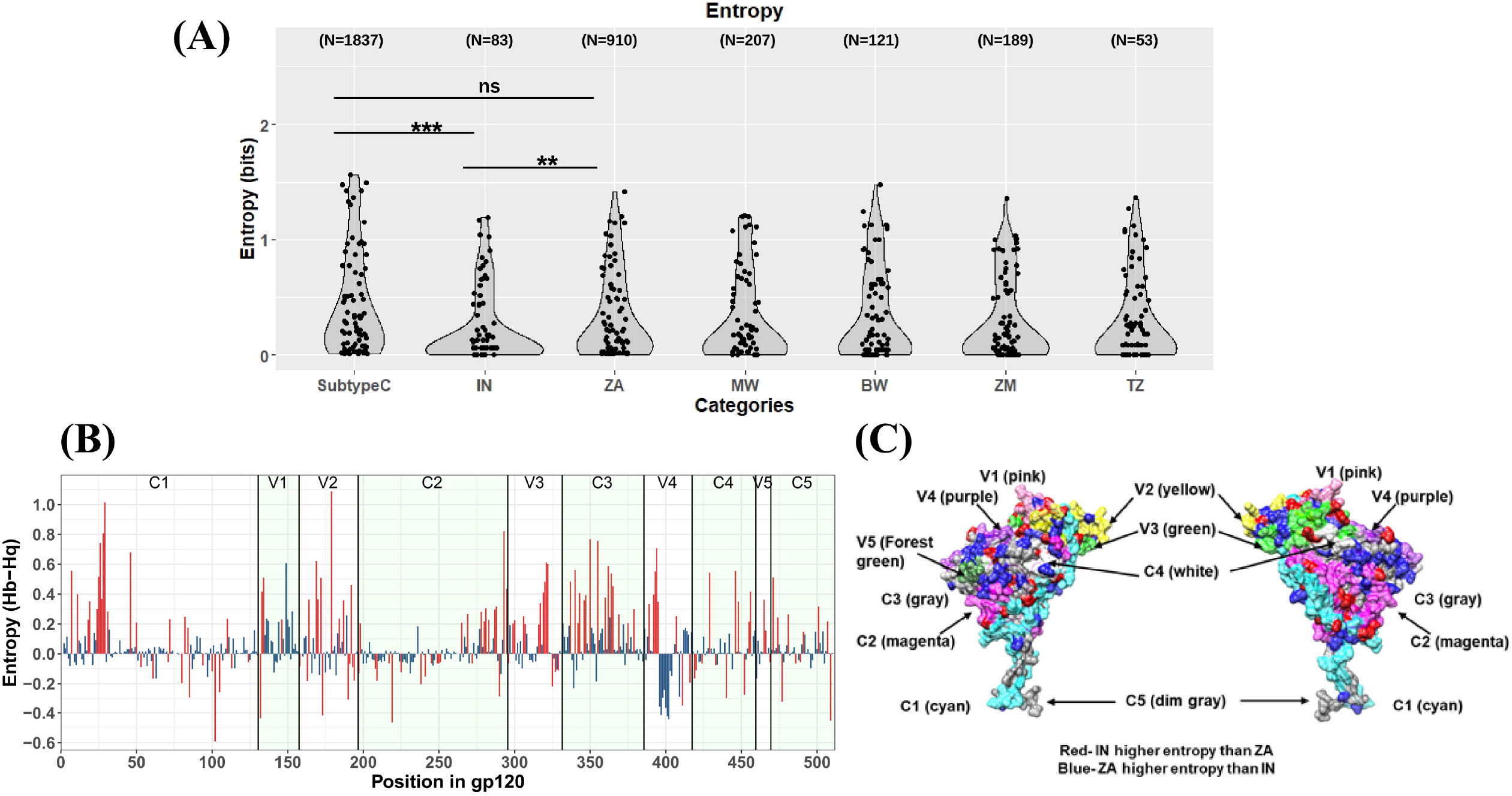
Entropy analysis. **A.** Shannon Entropy difference (H(background)-H(query) (unit: bits) has been plotted on Y axis against each amino acid position on X axis, where ZA dataset was the background while IN dataset was the query. Different domains of gp120 have been indicated. Bars with red color indicate positions with statistically significant entropy differences. Bars above 0 indicate higher entropy in South Africa while those below 0 indicate higher entropy in India. **B.** Variable entropy positions are plotted on prefusion gp120 envelope model derived from PDB:5U7O. gp120 domains have been color coded. Residue position highlighted in red indicate statistically significantly higher entropy in India compared to South Africa while those highlighted in blue indicate statistically significantly higher entropy in South Africa compared to India **(C)** Shannon entropy (bits) at key sites for bnAbs VRC01, VRC03, VRC07, VRC13, CAP256:VRC26.25, PGDM1400, PG9, PG16, PGT121 and PGT128 excluding positions in hypervariable regions have been plotted for overall Subtype C, South Africa (ZA) and India (IN). Statistical comparison p values have been obtained following application of Mann-Whitney test.

### Variation in Shannon entropy indicate significant intra-subtype C diversity

Shannon entropy is a measure of variability wherein higher entropy indicates higher variability. Indian sequence ‘query’ data set (N=83) was compared against South African sequence ‘background’ data set (N=910) through Entropy-TWO tool on LANL HIV database which generated Shannon entropy values with statistical confidence measured through Monte-Carlo randomization with 100 replacements. Overall, 133 amino acid positions across gp120 were detected to have differential entropy between India and South Africa, of which 83 sites had higher entropy in South Africa while 50 sites had higher entropy in India (Figure 4B). As indicated in Figure 4B, many of the differential entropy sites are located in the surface accessible V1-V2 and V3 regions, also targeted by several bnAbs. To assess the entropy differences associated with bnAb contact sites, we plotted entropy data for South Africa, India and HIV-1 subtype C overall, as reported in the HIV-LANL database. To prevent bias because of hypervariable nature of certain gp120 domains, the residue positions present in hypervariable regions were removed from the subsequent comparisons. While no difference was observed in entropy profiles at key bnAb sites between overall subtype C entropy and that observed in South Africa (Mann-Whitney test, p = 0.0759), corresponding entropy profile in India was significantly different both from overall subtype C (p<0.0001) and South Africa (p<0.0001). These observations suggest a differential pattern of conservation across the two populations and thus may result in variable breadth of neutralization by these bnAbs (Bai et al. 2019).

### Variation in abundances of epitopes associated with resistance and susceptibility to bnAbs

Next, we examined the abundance of HIV-1 subtype C resistance phenotype as defined in the CATNAP database of key bnAbs having varied epitope specificities in *gp120* across different countries. We took sequence datasets of different counties (www.hiv.lanl.gov) and calculated the frequency of key amino acid residues associated with the sensitivity and resistance to individual bnAbs. We examined the select key bnAbs targeting CD4bs (VRC01, VRC07, 3BNC117 and N6), V1V2 region (PG9, PG16, PGT145, PGDM1400 and CAP256.VRC26.25) and V3 supersite (PGT121, PGT128 and 10-1074). As shown in Figure 5, we found considerable variation in the abundance of amino acid residues that form epitopes associated with neutralization resistance to bnAbs with unique specificity. Variation in abundances of the following residues that form epitopes to bnAbs with unique specificities were observed in geographically divergent globally circulating HIV-1 subtype C (Figure 5). For CD4bs directed bnAbs: variation in the sensitivity to global HIV-1 subtype C to VRC01 was found to be majorly associated with N234 and S364. Similarly, for V1V2 directed bnAbs: K130, I161, Q170, Y173, S291, T297, N332 and N340 associated with significant variation in PGT145 sensitivity; K130, I161,I165, V169 and N332 (most significant) associated with significant variation in PGDM1400 sensitivity; K130, I161, R166, V169, Q170 and N332 (most significant) associated with significant variation in PG9 sensitivity; K130, I161 and K171 associated with significant variation in PG16 sensitivity; I165. R166, V169 and N332 associated with significant variation in CAP256.VRC26.25 sensitivity was observed. Finally, for V3 directed bnAbs: K155, I165, R252, N289, E293, I307, H330, N332, S334, A336 and N448 associated with significant variation in PGT121 sensitivity; R252, N295, N300, H330, S334, A336, Q344, and T415 associated with significant variation in PGT128 sensitivity; K155, N156, I165, N230, T240, R252, K282, I307, A316, H330, S334, A336, Q344 and N448 associated with significant variation in 10-1074 sensitivity were observed. Taken together, our observation indicates the existence of variation in the abundance of key bnAb contact sites across global circulating HIV-1 subtype C which further highlights that majority of the bnAbs likely would be effective when administered in proper combination against diverse region-specific circulating HIV-1 subtype C.

**Figure 5:**
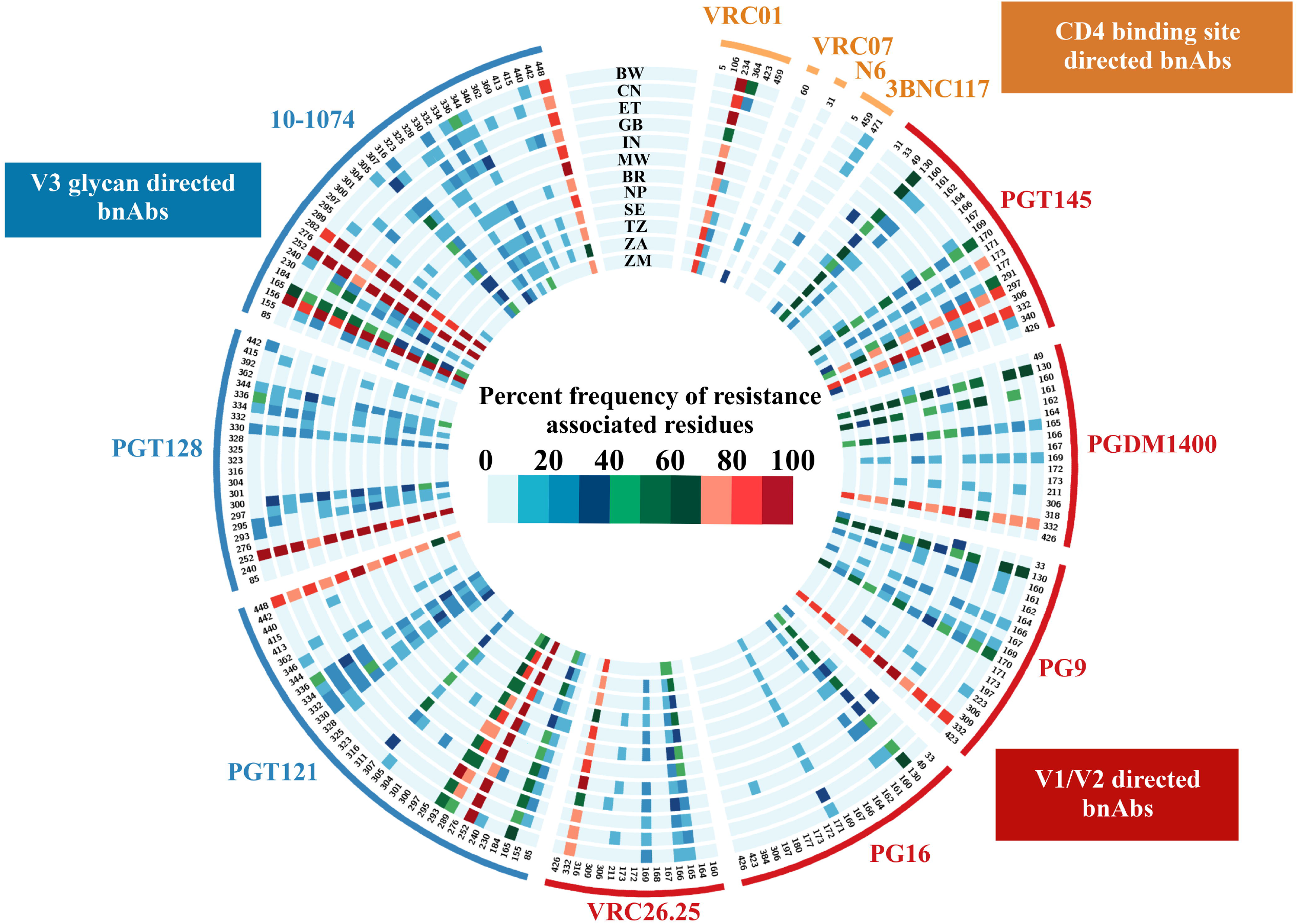
Abundance of bnAb resistance associated residues. A circos heatmap depicting abundance of bnAb resistance associated residues was plotted for 11 bnAbs (VRC01, VRC07, PGT121, PGT128, PGT145, PG9, PG16, VRC26.25, 3BNC117, 10-1074 and N6) wherein each track indicates the country of origin. Each pixel on the circular track indicates a specific residue position colored as per abundance of resistance causing residues at that position as per the color key.

### Evidence of accumulation of bnAb resistance phenotype in globally circulating HIV-1 C over time

With abundance of bnAb neutralization data against specific envelope sequences obtained *in vitro* as well as through clinical trials, several machine learning-based algorithms are increasingly becoming available that can predict probable sensitivity to bnAbs on the basis of gp120 sequences. In the present study we employed one such recently published algorithm bNAb-ReP (Rawi et al. 2019) to predict sensitivity of 1466 gp120 sequences selected in the present study from countries India, China, South Africa, Malawi, Zambia and Tanzania to following bnAbs: 3BNC117, VRC01, VRC07, PGT145, CAP256:VRC26.25, PGDM1400, PG9, PG16, PGT121, PGT128 and 10-1074. The prediction data were plotted along with country-wise *in vitro* data available through CATNAP database for a total of 283 sequences as indicated in Table 2. As indicated in Figure 6, probability values greater than 0.5 point towards sensitivity to bnAbs while those lower than 0.5 indicate probable resistance. For VRC01, while CATNAP database indicated unequivocal sensitivity of majority of sequences from all 6 countries, bNAb-ReP predicted a significant fraction of sequences from African countries to be resistant. Predictions for VRC26.25, 10-1074, PGT121 and PGDM1400 matched those reported in the CATNAP database. Similar to VRC01, predictions for 3BNC117, PGT128, PG9, PG16 and VRC07 did not match the data from CATNAP and indicated probable resistance in many of the sequences from all 6 countries. To assess if bnAb sensitivity differed over time, prediction data for each of the 11 bnAbs was plotted against three periods based on the reporting date of the sequences (Figure 7). The three periods considered were 1986-2000 (N=244), 2001-2010 (N=1187) an 2011-2019 (N=333). Except for PGT121, PGT128 and 10-1074, all bnAbs showed significant decrease in sensitivity over time. Despite this decrease, most sequences were predicted to be susceptible to PGDM1400 and VRC26.25. However, susceptibility to 3BNC117, VRC01, VRC07, PGT145, PG9 as well as PG16 was predicted to have reduced significantly over the period of time assessed.

**Table 2:** Details of sequences analyzed, and their sources used for predication analysis of HIV-1 subtype C to different bnAbs.

| Country | Number of sequences |  |
| --- | --- | --- |
|  | CATNAP | LANL-HIV (bNAbs-ReP) |
| India | 13 | 83 |
| China | 7 | 24 |
| South Africa | 146 | 910 |
| Malawi | 62 | 207 |
| Zambia | 23 | 189 |
| Tanzania | 32 | 53 |
| Total | 283 | 1466 |

**Figure 6:**
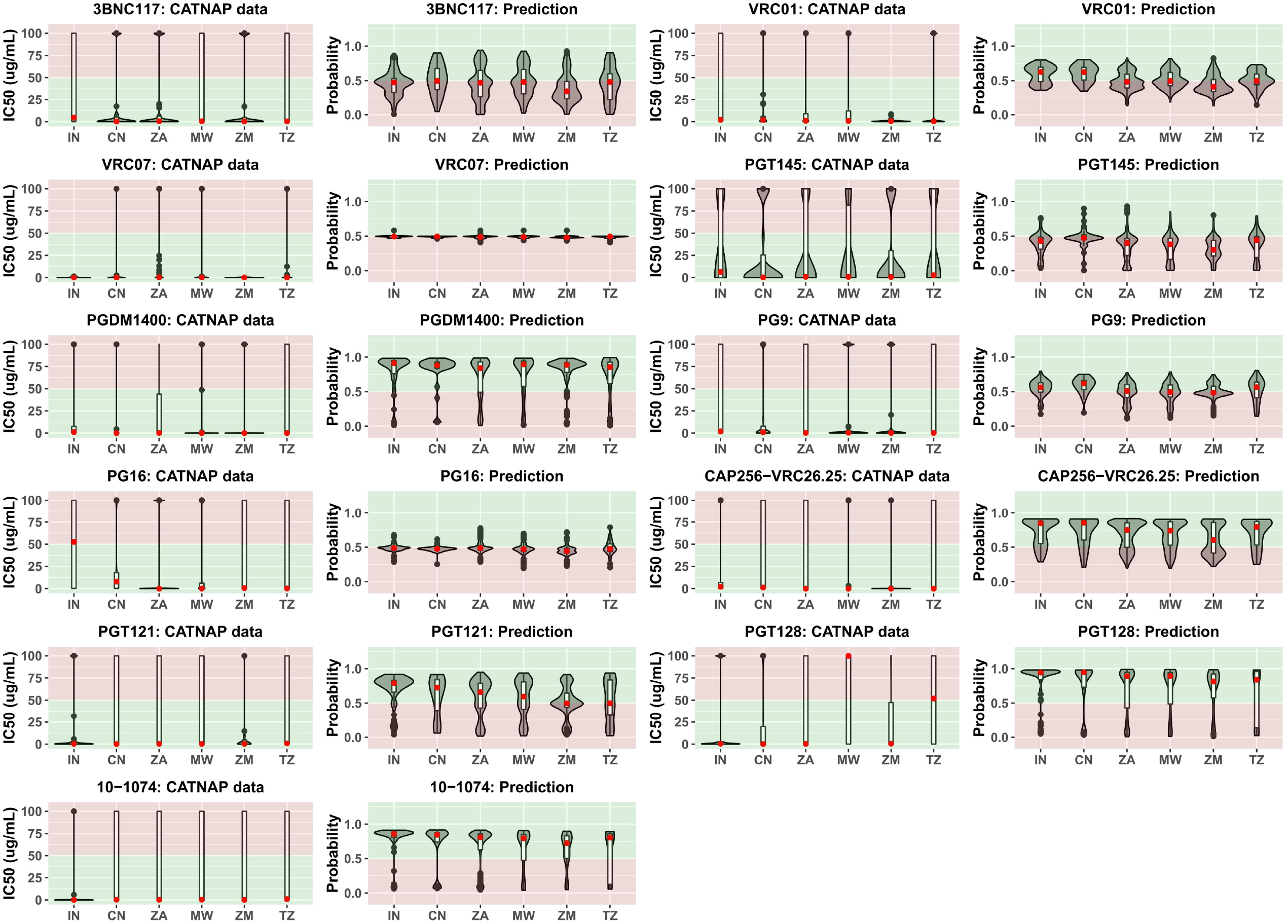
Prediction of bnAb sensitivity across different countries. Each panel indicates available country-wise CATNAP data plotted next to country-wise prediction data for available sequences for 3BNC117, VRC01, VRC03, VRC07, VRC13, CAP256:VRC26.25, PGDM1400, PG9, PG16, PGT145, PGT121, and PGT128 and 10-1074. In the CATNAP data panels, country-wise violin plots have been inlayed with boxplots against reported IC50 (μg/mL) values. Black dots indicate outliers while red dots indicate median values. Red background zone indicates bnAb resistance (IC50 >50 μg/mL) while green background zone indicates bnAb sensitivity (IC50 < 50 μg/mL). In the bNAb-ReP data panels, country-wise violin plots have been inlayed with boxplots against probability of neutralization as predicted by bNAb-ReP. Black dots indicate outliers while red dots indicate median values. Red background zone indicates probable bnAb resistance (Neutralization probability < 0.5) while green background zone indicates probable bnAb sensitivity (Neutralization probability > 0.5).

**Figure 7:**
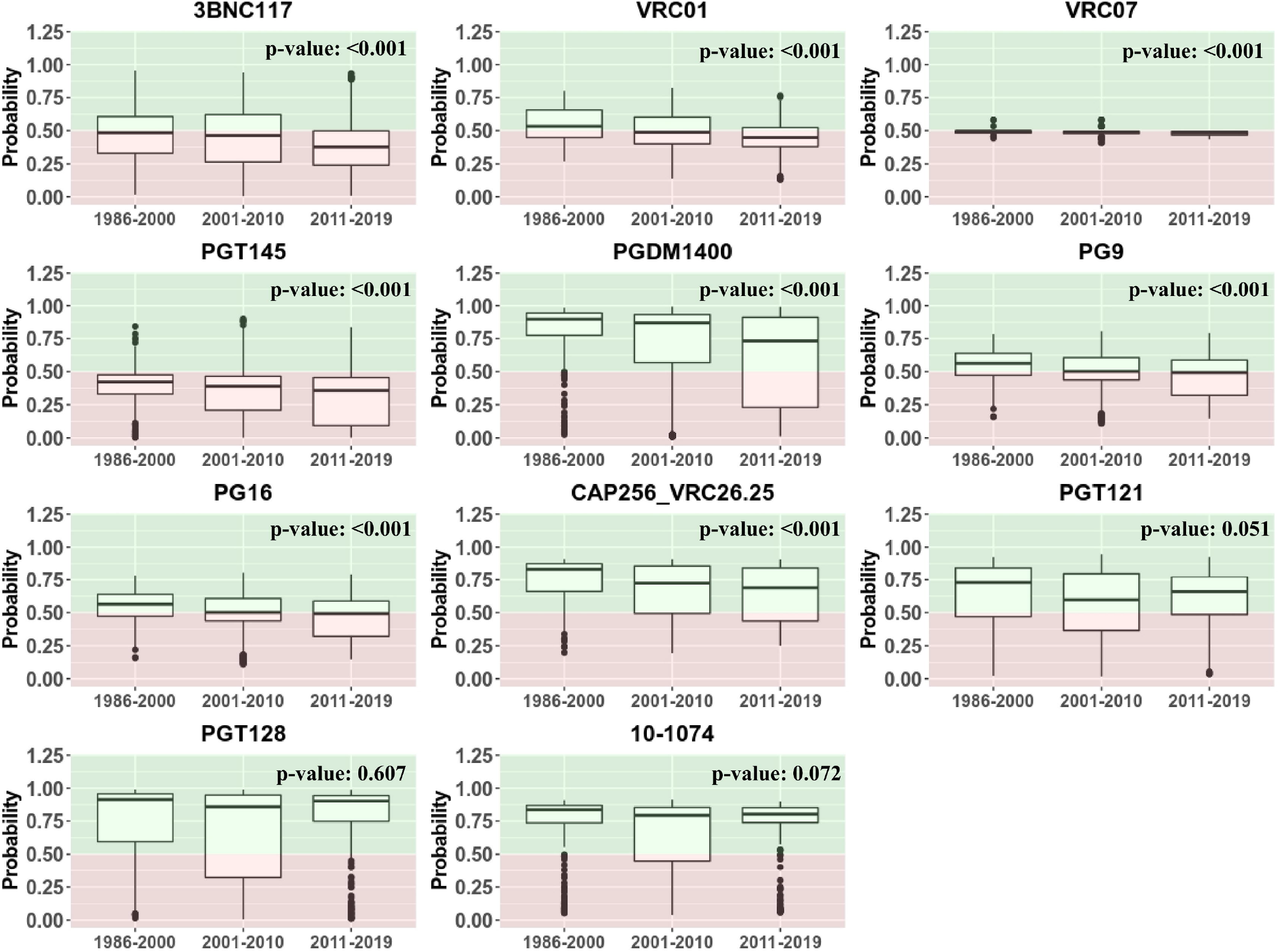
Assessment of predicted bnAb sensitivity over time. Each panel indicates cumulative prediction data for available sequences for 3BNC117, VRC01, VRC03, VRC07, VRC13, CAP256:VRC26.25, PGDM1400, PG9, PG16, PGT145, PGT121, and PGT128 and 10-1074 plotted against 3-time periods as follows: 1986-2000 (N=244), 2001-2010 (N=1187) and 2011-2019 (N=333). P values indicate trend analysis performed by Jonckheere-Terpstra test.

## Discussion

Given that subtype C accounts for approximately half of the global HIV burden, limited information on the intra-clade C *env* diversity and its association with variation in their neutralization phenotypes exists. In the present study, we compared the genetic attributes of the globally circulating HIV-1 subtype C *gp120* sequences that differentiate the region-specific intra-clade HIV-1 subtype C neutralization diversity. For this, we used the available information in the HIV database (www.hiv.lanl.gov) and established algorithm (Rawi et al. 2019) to suitably predict the association between genetic features that potentially dissects the HIV-1 intra-clade C neutralization diversity. Although there is a disparity between the number of existing region-specific unique HIV-1 subtype C sequences in the database, nonetheless, region-specific distinct genetic clustering was observed by phylogenetic analysis of *gp120* amino acid sequences. It is to be noted that our analysis was based on one sequence per individual to avoid any sampling bias on an individual level in our analysis. The region-specific subtype C *gp120* divergence could possibly be due to diversity in the population level across geography (Hraber et al. 2014). Indeed, studies have shown that HIV-1 can selectively incorporate broad range of biologically active host proteins in the process of viral egress, which can potentially exhibit altered pathogenicity and neutralization phenotypes (Burnie and Guzzo 2019). This also indicates possible association between intra clade C genetic diversity with ethnically distinct population.

The evolutionary genetic drift within subtype C that we observed from phylogenetic analysis is likely due to differential host characteristics which include immune response and differential genetic bottlenecks. For example, *gp120* sequences from Asian countries were found to demonstrate monophylectic clustering compared to that observed with those obtained from African countries. A number of studies (Deshpande et al. 2016; Ringe et al. 2012; van Gils et al. 2011) have demonstrated the role of loop length in the hypervariable regions, (with particular reference to V1V2 region), charge and N-linked glycosylation on altered neutralization phenotype. In the present study, we observed that while there is an existence of region-specific variation in V1V2 loop length, V4 loop length variation between subtype C sequences was found to be most profound across geographic boundaries. In addition, we also observed that the average *gp120* length of African countries was found to be smaller compared to other regions. Our data demonstrates subtle but significant variations in these attributes thereby indicating that a common set of bnAbs are not likely to be equally effective against the HIV-1 subtype C circulating globally. The above conclusion was further substantiated by our observation where we found significant variation in the entropy profiles (variation in sites/positions associated with different bnAbs) between the geographically distinct *gp120* sequences. With limited data available in CATNAP, indeed we found evidence of variation in susceptibility region-specific clade C to different bnAbs, which substantiate our observation. Interestingly, as reported elsewhere (Bouvin-Pley et al. 2013; Bouvin-Pley et al. 2014; Hake and Pfeifer 2017), our data also predicted potential likelihood of accumulation of resistance phenotype overtime to existing bnAbs. This observation indicate that it is necessary for continuous surveillance of evolving viruses in the context of subtype C along towards prioritizing bnAb combinations that will optimally dissect and overcome the evolving genetic diversity. Interestingly, a similar accumulation in ART resistance has been documented and studies that concurrently evaluate the interplay of these two evolutionary patterns, heretofore considered to be mutually exclusive, may highlight novel and synergistic therapeutic strategies. In summary, the differences in HIV-1 clade C *gp120* sequences observed herein indicate disparate and distinctly evolving clusters within clade C with differential predicted responses to bnAbs. Elucidation of neutralization diversity of subtype C particularly in context of evolution of *gp120* overtime will be essential for selecting appropriate bnAb combination for effective prophylaxis and treatment and also in informing rational vaccine design. In summary, our study highlights that towards developing HIV-1 bnAbs as products for prevention and treatment, continued surveillance of the evolution of genetic features with particular reference to *env* gene that are targets of neutralizing antibodies of globally circulating HIV-1 remains crucial for the identification and prioritization of combination of bnAbs that would best suit to provide maximal geography and population-specific neutralization coverage.

## Materials and Methods

### Retrieval of gp120 sequences

Sequences for the gp120 gene were retrieved from manually curated LANL HIV database (www.hiv.lanl.gov). Briefly, HIV-1 subtype C nucleotide sequences fully covering the genomic region 6225-7758 (as per HXB2 numbering) were retrieved and subsequently filtered with a one sequence per individual filter criterion. Sequence entries without any information regarding the sample source country were excluded. Multiple sequence alignment for the amino acid sequences along with HXB2 sequence (GenBank: K03455.1) was produced with Gene cutter (“Gene Cutter,” LANL). Gene Cutter clips the coding regions from unaligned nucleotide sequences and produces amino acid alignments based on Hmmer v 2.32 algorithm with a training set of the full-length genome alignment. Alignments were manually curated using Bioedit v7.2.5 (Hall 1999). Sequences with internal stop codons were discarded.

### Phylogenetic analysis

The number of amino acid differences per site were estimated by averaging over all sequence pairs between different countries using Molecular Evolutionary Genetics Analysis software (MEGA v.10) (S. Kumar et al. 2018b). The rate variation among sites was modelled with a gamma distribution (shape parameter = 1). This analysis involved 1814 amino acid sequences. Phylogenetic trees were generated for the amino acid alignments with iqtree under ‘HIVb’ model with estimated Ƴ parameters and number of invariable sites (Nguyen et al. 2015). Robustness of the tree topology was further assessed by SH-aLRT as well as 1000 ultrafast bootstrap replicates implemented in iqtree. A subtree consisting of 251 sequences were again constructed as mentioned previously.

### Estimation of the variable loop properties and potential N linked glycosylation sites

Variable loop regions for V1 (131-157: HXB2 numbering), V2 (158-196), V3 (296-331), V4 (386-417) and V5 (460-469) were retrieved from amino acid alignments with Bioedit v7.2.5. Each of the loop datasets were then processed with custom bash/awk scripts to generate length statistics. The length distributions were further assessed and compared by Kruskal Wallis test followed by Dunn’s multiple comparison test. Cumulative variable loop charge values were predicted for each of the sequences with custom bash scripts wherein, Lysine (K), Arginine (R) and Histidine (H) residues were assigned +1 values each while Aspartic acid (D) and Glutamic acid (E) were assigned −1 values each. Potential N-linked glycosylation sites were predicted in amino acid sequence datasets with N-GlycoSite tool hosted at the HIV-LANL database (Zhang et al. 2004). Prevalence of each of the pNLG sites under study were calculated using custom bash/awk scripts and were further assessed statistically using Fisher’s exact test. Country-wise pNLGs abundance heatmap was plotted using ‘pheatmap’ package in R.

### Entropy analysis

Shannon entropy for selected sequence data sets was generated using Entropy-two tool following 100 randomizations with replacement (Efron and Tibshirani 1991; Gaschen et al. 2002) (www.hiv.lanl.gov. Key sites for interaction with bnAbs VRC01, VRC03, VRC07, VRC13, CAP256:VRC26.25, PGDM1400, PG9, PG16, PGT121 and PGT128 were derived from CATNAP database. HIV-1 clade C overall entropy bed graph was derived from Genome browser on HIV-LANL database.

### Analysis of bnAb epitope contact sites

Specific epitope contact sites/positions along with documented variants imparting sensitivity or resistance phenotype were retrieved from CATNAP database for bnAbs: VRC01, VRC07, PGT121, PGT128, PGT145, PG9, PG16, VRC26.25, 3BNC117, 10-1074 and N6. Frequency of such resistant variants were calculated across sequences from countries with more than 20 sequences available, using in-house bash scripts. Contact site resistance heatmaps were prepared with Circos v0.61 (Krzywinski et al. 2009).

### Prediction of bnAb sensitivity with bNAb-ReP

Sensitivity of contact sites to bnAbs (3BNC117, VRC01, VRC07, CAP256:VRC26.25, PGDM1400, PG9, PG16, PGT145, PGT121, PGT128 and 10-1074) were predicted with bNAb-ReP tool (Rawi et al. 2019). Neutralization data corresponding to the selected bnAbs was retrieved from CATNAP database (www.hiv.lanl.gov).

The temporal prediction data was stratified into approximately three decades as follows: 1986-2000 (N=244), 2001-2010 (N=1187) and 2011-2019 (N=333).

### Statistical analyses and data presentation

Statistical analyses for variable loop length distributions were performed using GraphPad Prism version 5.01 for Windows, GraphPad Software, San Diego California USA. Statistical Comparison of Fisher’s test for pNLG sites as well as abundance of bnAb resistance associated residues was performed through R statistical computing software (v3.4.0) and R studio v1.0.143 (R. Team 2015; R. C. Team 2018). Phylogenetic trees were visualised and edited with the R package ‘Graphlan’ (Asnicar et al. 2015). Variable entropy positions were plotted on prefusion gp120 envelope model derived from PDB:5U7O in Chimera v1.14 (Pettersen et al. 2004). Plots depicting variable region characteristics, entropy differences and bNAb sensitivity predictions were prepared using ‘ggplot2’ package in R (Analysis.. 2016). Trend analysis for predicted bnAb sensitivity was performed by Jonckheere-Terpstra test implemented in R statistical software.

## Acknowledgements

We thank our laboratory members for providing valuable inputs and suggestions. The authors wish to acknowledge the funding support from the Wellcome Trust/DBT India Alliance Team Science Grant (IA/TSG/19/1/600019), Science & Engineering Research Board, Department of Science & Technology, Government of India (CRG/2019/002939) and Department of Biotechnology, Government of India (BT/PR24520/MED/29/1222/2017). ICMR-NIRRH is acknowledged for server support for computation analysis. IAVI’s work was made possible by generous support from many donors, including the Bill & Melinda Gates Foundation, the Ministry of Foreign Affairs of Denmark, Irish Aid, the Ministry of Finance of Japan, the Ministry of Foreign Affairs of the Netherlands, the Norwegian Agency for Development Cooperation (NORAD), the United Kingdom Department for International Development (DFID), and the United States Agency for International Development (USAID). The full list of IAVI donors is available at www.iavi.org. The contents are the responsibility of the International AIDS Vaccine Initiative and do not necessarily reflect the views of USAID or the United States Government. We thank Prof Gagandeep Kang, Translational Health Science & Technology Institute for support.

## Notes

### Competing Interest Statement

The authors have declared no competing interest.

### Summary of Updates

Geospatial HIV-1 subtype C gp120 sequence diversity and its predicted impact on broadly neutralizing antibody sensitivity

